# Commensal *Pseudomonas fluorescens* protect *Arabidopsis* from closely-related *Pseudomonas* pathogens in a colonization-dependent manner

**DOI:** 10.1101/2021.09.02.458786

**Authors:** Nicole R. Wang, Ryan A. Melnyk, Christina L. Wiesmann, Sarzana S. Hossain, Myoung-Hwan Chi, Kitoosepe Martens, Cara H. Haney

## Abstract

Plants form commensal associations with soil microorganisms, creating a root microbiome that provides benefits to the host including protection against pathogens. While bacteria can inhibit pathogens through production of antimicrobial compounds *in vitro*, it is largely unknown how microbiota contribute to pathogen protection *in planta*. We developed a gnotobiotic model system consisting of *Arabidopsis thaliana*, and an opportunistic pathogen *Pseudomonas* sp. N2C3, to identify mechanisms that determine the outcome of plant-pathogen-microbiome interactions in the rhizosphere. We screened 25 phylogenetically diverse *Pseudomonas* strains for their ability to protect against N2C3 and found that commensal strains closely related to N2C3 were more likely to protect against pathogenesis. We used a comparative genomics approach to identify unique genes in the protective strains that revealed no genes that correlate with protection, suggesting that variable regulation of components of the core *Pseudomonas* genome may contribute to pathogen protection. We found that commensal colonization level was highly predictive of protection and so tested deletions in genes previously shown to be required for *Arabidopsis* rhizosphere colonization. We identified a response regulator *colR* that is required for *Pseudomonas* protection from N2C3 and fitness in competition with N2C3 indicating that competitive exclusion may contribute to pathogen protection. We found that *Pseudomonas* WCS365 also protects against the agricultural pathogen *Pseudomonas fuscovaginae* SE-1, the causal agent of bacterial sheath brown rot of rice. This work establishes a gnotobiotic model to uncover mechanisms by which members of the microbiome can protect hosts from pathogens and informs our understanding of the use of beneficial strains for microbiome engineering in dysbiotic soil systems.

## Introduction

Plants exist in collaboration with microorganisms that colonize their tissues, forming microbiomes in ecological niches such as the phyllosphere (leaf surface) or rhizosphere (root surface). Due to the diverse collection of genes provided by each member of the rhizosphere microbiome, the rhizosphere microbiome has great genetic potential to beneficially impact the plant host. Plants are able to harness the genetic potential of the rhizosphere microbiome by shaping the community composition in response to changes in environmental conditions [1,2]. The presence of specific beneficial bacteria, in turn, can protect against pathogens either directly, through antibiosis, or indirectly, by priming the plant immune system for future attack or by outcompeting pathogens for space and nutrients [3]. However, the genes and mechanisms that govern these beneficial interactions are not fully understood [4].

Associations between *Arabidopsis* and *Pseudomonas* species can serve as model systems to study plant-microbe interactions in the rhizosphere [5,6]. *Pseudomonas fluorescens* and related species are widely plant-associated and are ubiquitously present in the rhizosphere microbiome. *Pseudomonas* are enriched in disease suppressive soils [7] and are known to produce diverse antimicrobial compounds [8]. The inoculation of single strains of *P. fluorescens* is sufficient to provide benefits to the plant, such as promoting root growth, or acting as a biocontrol agent, making *Pseudomonas* strains useful in a reductionist approach to studying beneficial plant-microbe interactions [6].

The species complex *P. fluorescens* is genetically and functionally diverse, despite being over 97% identical based on 16S rRNA gene sequence [9,10]. Even within closely-related *P. fluorescens* strains, host-associated lifestyles can differ. For example, the Brassicacearum clade of *Pseudomonas* contains the pathogenic *Pseudomonas* sp. N2C3 amongst numerous plant growth-promoting bacteria such as *Pseudomonas* sp. WCS365 and *P*. *brassicacearum* NFM421 [9]. *Pseudomonas* sp. N2C3 can cause disease and stunting on plants within the Brassicaceae (including *Arabidopsis*, broccoli, kale) and Papaveroideae (poppy) families when grown in gnotobiotic conditions [9]. However, when inoculated in natural soil, N2C3 was unable to stunt *Arabidopsis* growth [2]. This suggests that the presence of beneficial microbes in the rhizosphere may lead to protection against N2C3 pathogenesis.

Using *Pseudomonas* sp. N2C3 pathogenesis of *Arabidopsis* as a model system, we screened genome-sequenced commensal *Pseudomonas* for protection against this opportunistic pathogen. Closely-related *Pseudomonas* strains with markers of both commensal and pathogenic lifestyles have been identified within the microbiome of the same plant [9,11] indicating they naturally co-occur. We identified a number of *Pseudomonas* strains that are close relatives of N2C3 that can protect against pathogenesis and used this model to identify genetic mechanisms that shape plant-pathogen-microbiome interactions.

## Results

### Closely-related *Pseudomonas* strains protect from an opportunistic pathogen

We previously observed that the *Pseudomonas* pathogen N2C3 readily causes disease under gnotobiotic conditions [9], but fails to cause disease in soil [2], suggesting that members of the microbiome may protect against pathogenesis. To test this, we used *Pseudomonas* sp. N2C3 and *Arabidopsis* as a model to identify bacterial strains that are protective against N2C3 pathogenesis in the rhizosphere under gnotobiotic conditions. As the *Pseudomonas* genus is composed of genetically diverse species that are often present in the rhizosphere microbiome, 25 strains spanning the diversity of the *Pseudomonas* genus were tested for their ability to protect against N2C3 (Table 1). We identified 10 *Pseudomonas* strains that when co-inoculated at a 5:1 ratio with N2C3 increased the fresh weight of *Arabidopsis* (Fig 1A). We also identified 11 strains that failed to protect against N2C3 as indicated by a similar fresh weight to seedlings treated with N2C3 alone. An additional 4 strains stunted plants, including two *P. aeruginosa* strains known to be pathogenic on plants [12,13], and two *P. fluorescens-clade* stains that have a pathogenicity island encoding syringomycin and syringopeptin [9], indicating they are pathogenic. Collectively this indicates that commensal *Pseudomonas* can protect from an opportunistic pathogen under gnotobiotic conditions and that there is natural variation in the protective ability across the genus *Pseudomonas*.

**Table 1.**
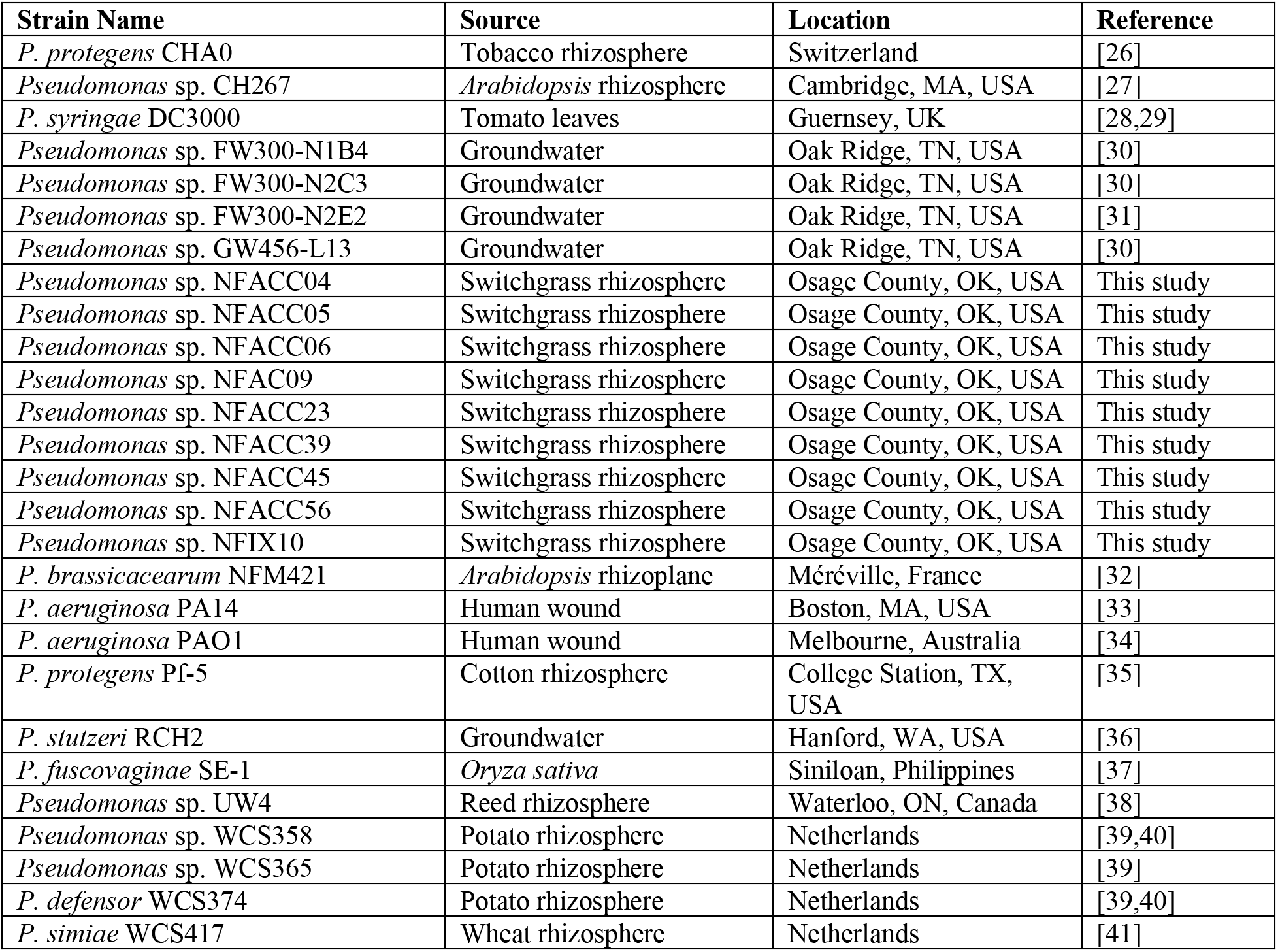
Strains used in this study.

**Figure 1.**
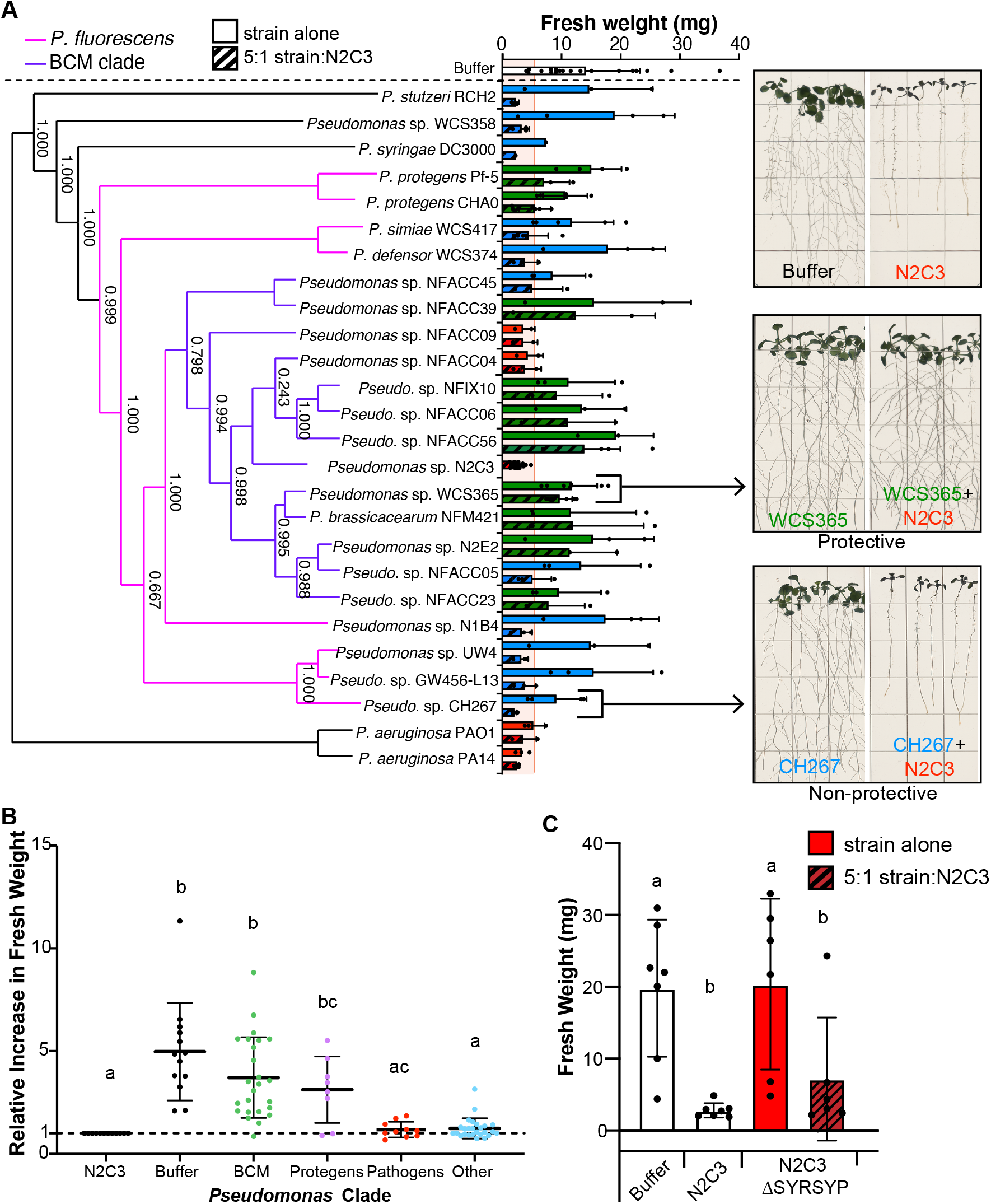
Phylogenetically related commensal *Pseudomonas* can protect against the *Pseudomonas* pathogen N2C3. (A) The ability to protect against N2C3 (green bars) is prevalent within the Brassicacearum/Corrugata/Mediterranea (BCM) clade (purple). Beneficial *Pseudomonas* strains outside of the BCM clade, with the exception of Protegens strains Pf-5 and CHAO, are unable to protect (blue). *Pseudomonas* strains were co-inoculated with N2C3 at a 5:1 ratio. Fresh weight of *Arabidopsis* seedlings was used as a proxy for protection against N2C3. Pathogenic strains that stunt plant growth alone are marked in red. Dashed line represents the mean + 3 standard deviations of N2C3-treated plants, estimating where 99.7% of measurements should fall under a normal distribution. Means that surpass this threshold were classified as protective (green bars). Representative images of buffer, pathogen (N2C3), a protective strain (WCS365) and a non-protective strain (CH267) are shown. (B) Strains within the BCM and Protegens clades significantly increase plant fresh weight when co-inoculated with N2C3, while non-pathogenic strains in other clades do not. Plant fresh weights were normalized by dividing by the average fresh weight of N2C3-treated control plants for the experiment. (C) Avirulent *Pseudomonas* sp. N2C3 cannot protect against virulent N2C3. A strain with a deletion of the pathogenicity island encoding syringomycin and syringopeptin was added at a 5:1 ratio with virulent N2C3. (A-C) Each data point represents an average of 4-5 plants co-inoculated with the test *Pseudomonas* strain and N2C3 from a single experiment. All repeated treatments tested in independent experiments were included as separate data points. Statistics were calculated using a one-way ANOVA and Tukey’s HSD. Lines represent mean +/-SD.

We noted that 8 of the 10 protective *Pseudomonas* strains are within the Brassicacearum, Corrugata, and Mediterranea (BCM) clade [9] and the other two are within the Protegens clade (Fig 1A) suggesting there may be some phylogenetic groups of *Pseudomonas* that are more able to protect against N2C3 pathogenesis than others. Using the data shown in Figure 1A, we tested whether strains in the BCM and Protegens clade were significantly more likely to protect than other strains. We found a significant increase in *Arabidopsis* weights when seedlings were treated with strains from either the BCM or Protegens clades, but not other commensal *Pseudomonas* strains (Fig 1B) supporting that these phylogenetic groups are more likely to be protective against N2C3.

Because of the strong phylogenetic signature of protection within the BCM clade, we wondered if an avirulent *Pseudomonas* sp. N2C3 mutant (a BCM strain) would be able to protect against virulent wild-type N2C3. N2C3 causes disease through quorum signaling-dependent synthesis of the non-ribosomal peptides syringomycin (SYR) and syringopeptin (SYP) [9]. We tested an avirulent N2C3 mutant with a deletion of SYR/SYP for its ability to protect against the wildtype N2C3. Consistent with previous findings, the ΔSYR/SYP mutant was no longer pathogenic (Fig 1C [9]); however, it was unable to protect against pathogenic N2C3 (Fig 1C). These data suggest that mechanisms responsible for protection against N2C3 are uniquely present in the non-pathogenic BCM strains.

### Gain and loss of components of the *Pseudomonas* accessory genome does not explain protection against *Pseudomonas* sp. N2C3

A number of phenotypes within *Pseudomonas*, ranging from benefits to plants [9,10], and virulence on plants and animals [9,14], can be attributed to gain and loss of components of the *Pseudomonas* accessory genome. To identify genes that may correlate with protection, we used PyParanoid, a previously described comparative genomics platform [9], to identify genes that were unique to protective strains. To test whether Pf-5 obtained genes responsible for protection in the BCM clade through horizontal gene transfer, we identified 14 genes that were present in 7 BCM strains (N2E2, NFACC23, NFACC39, NFACC45, NFIX10, and WCS365) and Pf-5, but absent in 7 non-protective strains outside of the BCM clade (CH267, GW456-L13, N1B4, UW4, WCS358, WCS374, and WCS417) (Figure S3A). Both protective and non-protective BCM strains were included in this comparison, to prevent exclusion of genes that are differentially expressed within the BCM clade. 12 out of 14 of these genes were deleted in WCS365, but none of them had a significant impact on its ability to protect against N2C3 (Figure S3B).

We then searched for genes present in seven protective BCM strains (NFM421, N2E2, WCS365, NFIX10, NFACC06, NFACC39, and NFACC56), the protective non-BCM strain Pf-5, but absent in N2C3. This approach yielded only 1 unique gene (group_03914), encoding a CinA family protein (Figure S4A). However, this gene was subsequently found to be present in 9 non-protective strains (NFACC05, NFACC45, N1B4, PA14, PAO1, RCH2, WCS358, WCS374, and WCS417) and absent in the protective strain CHA0, indicating that it is unlikely to underlie protection. These comparisons suggest that the Protegens and BCM clades may use distinct mechanisms for pathogen protection.

We repeated the comparative genomics analyses using just strains within the BCM clade, to identify if the presence any gene groups were responsible for differences in protection ability within this clade. To identify genes unique to protective BCM strains, we searched for genes present in 8 protective BCM strains (NFM421, N2E2, WCS365, NFIX10, NFACC23, NFAC45, NFACC39, NFACC06, and NFACC56) but absent in 3 non-protective BCM strains (NFACC45, NFACCO5, and N2C3). Using this approach, we identified no genes unique to the protective BCM strains (Fig S4B). To identify genes that are unique to non-protective BCM strains, we conducted the opposite analysis, searching for genes that were present in 3 non-protective BCM strains (NFACC45, NFACC05, and N2C3), but absent in 8 protective BCM strains (NFM421, N2E2, WCS365, NFIX10, NFACC23, NFAC45, NFACC39, NFACC06, and NFACC56). Using this approach, we identified 1 gene group encoding an aldo/keto reductase (group_06773) that was unique to non-protective BCM strains (Fig S5A). However, when the presence/absence of this gene group was plotted onto a tree spanning the diversity of the *Pseudomonas* genus, we observed that not all non-protective strains outside of the BCM clade possessed this gene group, indicating that the absence of this gene was unlikely to be responsible for protection in the BCM clade.

To identify genes whose absence may be responsible for a BCM clade-specific mechanism of protection, we searched for genes that were present in 9 non-BCM strains (Pf-5, CH267, GW456-L13, N1B4, RCH2, UW4, WCS358, WCS374, and WCS417) but absent in 7 protective BCM strains (N2E2, NFACC06, NFACC39, NFACC56, NFIX10, NFM421, and WCS365) (Fig S5B). However, we were unable to identify any genes that met this criterion. These data suggested that genes uniquely present or absent I n the protective or non-protective BCM clade members were not responsible for their ability to protect against N2C3.

A previous genome-wide association study identified three genomic regions that are negatively correlated with pathogenicity within the BCM clade, and are typically only present in the non-pathogenic members of the clade [9]. These genomic regions encode a type III secretion system (T3SS), diacetylphloroglucinol (DAPG) biosynthesis genes, and a T3SS effector, HopAA [9,15] (Fig 2A). To test whether T3SS or DAPG could contribute to protection against N2C3, we generated clean deletions of the *hrcC* gene [16] and DAPG biosynthesis island in the protective strain *Pseudomonas* sp. WCS365. We found that the *Pseudomonas* WCS365 *ΔhrcC* and ΔDAPG mutants had similar levels of protection as wild-type WCS365 (**Error! Reference source not found.**B). As we were unable to identify components of the *Pseudomonas* accessory genome involved in protecting against N2C3 we instead hypothesized that regulation of core *Pseudomonas* genetic components, rather than presence/absence of genes, may be required for protection.

**Figure 2.**
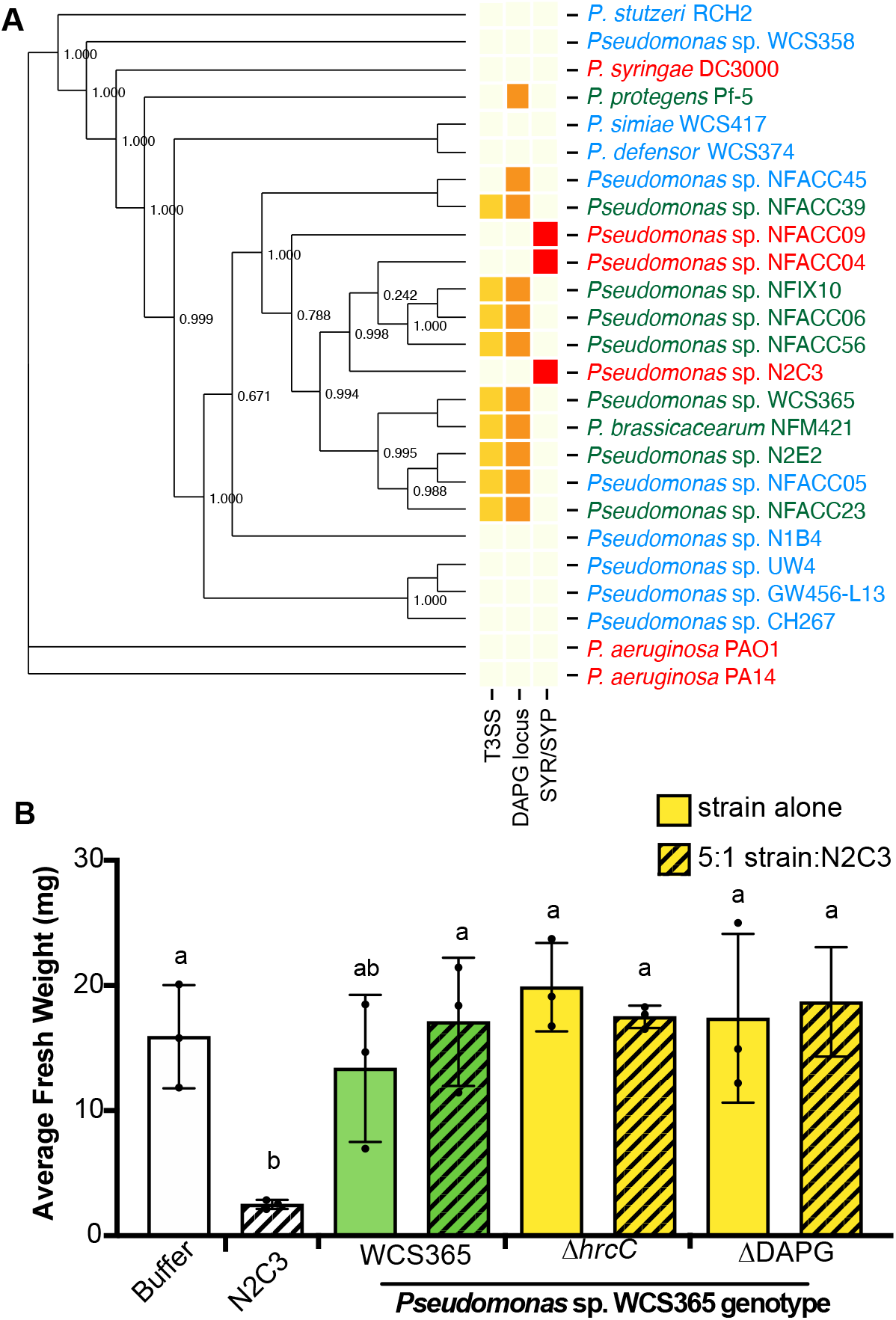
The ability to protect against N2C3 across the genus *Pseudomonas* is not due to horizontal transfer of components of the accessory genome. (A) Comparative genomics failed to identify any genes unique to protective strains (Figs S3 and S4). A previous GWAS identified a T3SS and diacetylphloroglucinol (DAPG) biosynthesis genes that are anti-correlated with pathogenicity in the BCM clade [7]. Strain names are colour coded according to their ability to protect (green), not protect (blue) or stunt plants alone (red). (B) *Pseudomonas* sp. WCS365 Δ*hrcC* (deficient in T3SS) and ΔDAPG mutants still protect against N2C3. Letters indicate p<0.05 as determined using a one-way ANOVA and Tukey’s HSD test. Each dot represents the average weight of 5 plants from a single biological replicate; separate dots are independent experiments with mean +/-SD plotted.

### Colonization ability predicts protection against *Pseudomonas* sp. N2C3

To gain insights into how protective strains may protect *Arabidopsis*, we first tested whether the protective *Pseudomonas* sp. WCS365 could directly kill N2C3 or inhibit its virulence. Using a bacterial cross streak assay, we found that WCS365 did not inhibit growth of N2C3 (Fig S1A). To test whether WCS365 can inhibit quorum signaling by N2C3, we used a *Chromobacterium violaceum* CV026 biosensor for C4-C8 AHL molecules, which reports AHLs from N2C3 through the production of the purple pigment violacein (Fig. S1B [9,17]). We found that violacein production was maintained when WCS365 and N2C3 were mixed and plated onto LB agar, indicating that WCS365 does not suppress N2C3 quorum signaling (Fig. S1B). These data indicate that WCS365 does not directly kill N2C3 nor does it interfere with regulation of SYR/SYP-production in N2C3 *in vitro*.

Since we found that *Pseudomonas* sp. WCS365 does not kill N2C3 or inhibit quorum signaling *in vitro*, we hypothesized that protective strains may outcompete the pathogen in the rhizosphere. To test this, we generated a N2C3-lacZ transposon insertion mutant to allow us to homogenize roots and the attached bacteria, and distinguish CFUs from distinct *Pseudomonas* strains on media containing X-gal. We found that protective strains (WCS365, N2E2 and Pf-5), maintained at least a 5:1 ratio (similar to the initial inoculum) in competition with N2C3-lacZ (Fig 3A). In contrast, CH267, which cannot protect *Arabidopsis* from N2C3, was significantly outcompeted by N2C3 (Fig 3A). We tested whether competition against N2C3 could predict protection in commensal *Pseudomonas* strains and found a non-linear relationship with a threshold for protection of approximately 5:1 (**Error! Reference source not found.**B; Fig S2). These data indicate that colonization may be a prerequisite for pathogen protection among *Pseudomonas* spp. strains.

**Figure 3.**
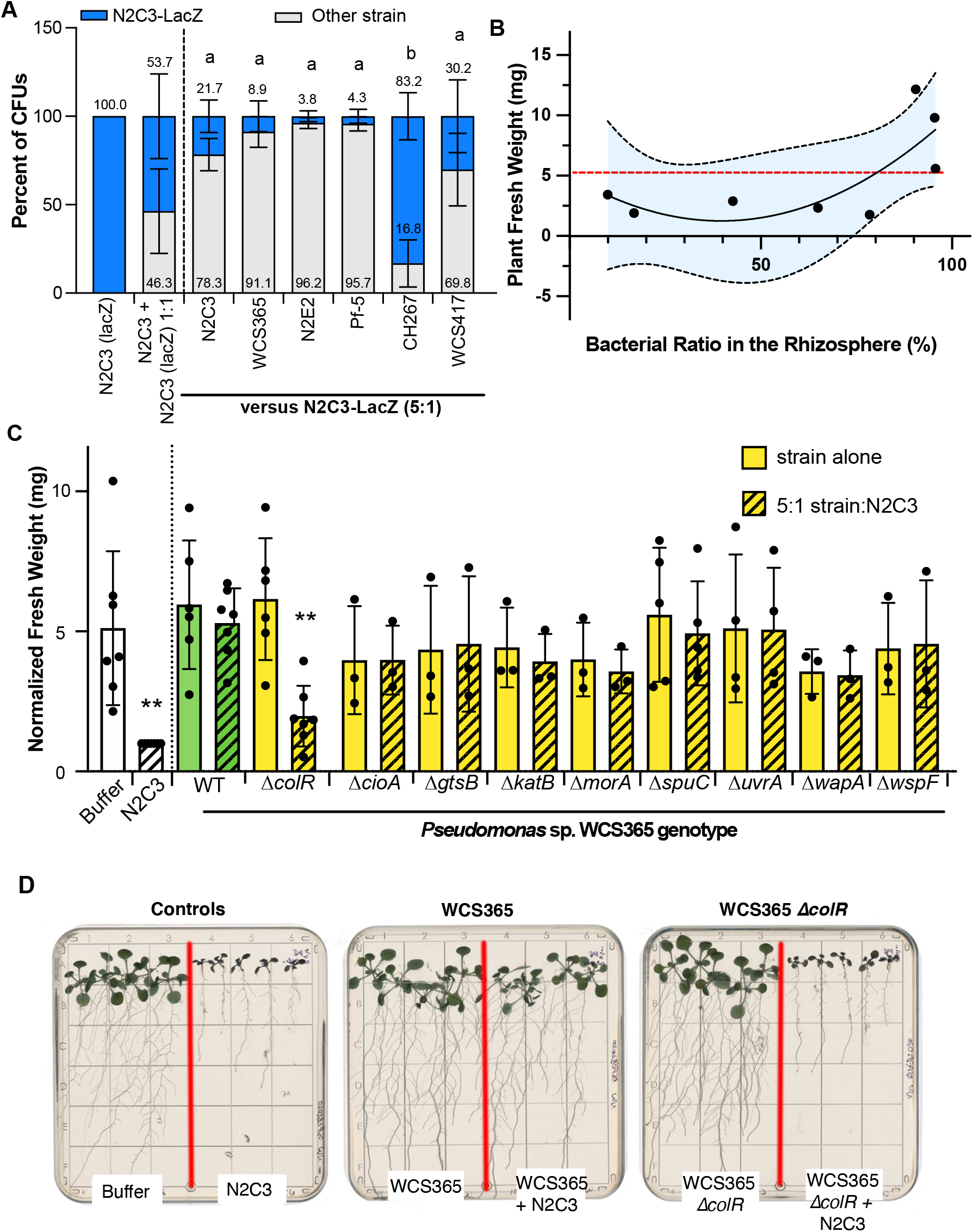
Protection against *Pseudomonas* sp. N2C3 is rhizosphere colonization dependent. (A) *Pseudomonas* strains that protected against N2C3, such as members of the BCM clade (WCS365, N2E2) and Pf-5, maintained at least a 5:1 ratio when in competition with N2C3, while strains that cannot protect had decreased fitness when co-inoculated with N2C3. *Pseudomonas* strains were competed against a N2C3 expressing *lacZ*. Relative abundance of each strain was determined by grinding roots, plating on media containing X-gal, and counting blue and white colonies. Letters denote significance (p<0.05) by two-way ANOVA and Tukey HSD. Bars represent mean +/-SD. (B) Correlation between the bacterial fitness in the rhizosphere and the plant fresh weight. Data points represent the averaged fresh weights and average ratios for the strain co-inoculated with N2C3-lacZ. Non-linear regression plotted with shaded area representing 95% confidence intervals. The red dashed line denotes the upper 99.7% confidence interval for increase in plant weight when co-inoculated with N2C3 as determined in Figure 1. (C) *Pseudomonas* sp. WCS365 deletion mutants of genes previously shown to be required for colonization of the *Arabidopsis* rhizosphere. Only deletion of the response regulator *colR* resulted in a loss in ability of *Pseudomonas* sp. WCS365 to protect *Arabidopsis* from N2C3. Each strain was co-inoculated onto roots with N2C3 at a 5:1 ratio. **p<0.05 relative to WT WCS365 + N2C3 determined by one-way ANOVA and Tukey HSD. Mean +/-SD is plotted. (D) Representative images showing loss of protection by *Pseudomonas* sp. WCS365 Δ*colR* mutant.

Since we found that colonization was correlated with protection against *Pseudomonas* sp. N2C3 (Fig 3B), we hypothesized that competitive exclusion could be important for protection. During competitive exclusion, existing members of the microbiome protect against pathogens by competing for space and nutrients rather than direct antagonism. To test whether protection from N2C3 is colonization dependent, we tested 8 WCS365 deletion mutants that were found to have fitness defects in a previous Tn-Seq screen [18] for their ability to protect against N2C3. We found that the majority of genes tested are broadly conserved across protective and non-protective strains (Fig S6). Of the mutants tested, we found that only WCS365 Δ*colR* lost its ability to protect against N2C3 (Fig 3C). ColR is a response regulator in a two-component system that has previously been shown to be necessary for colonization of plant roots [18,19]. This indicates that protection against N2C3 relies on a ColR-dependent mechanism for colonization.

ColR/S is a rhizosphere colonization or fitness determinant of *Pseudomonas* in both *Arabidopsis* and potato [18,19]. Several genes expressed in a ColR/S-dependent manner have been identified including several required for rhizosphere colonization. These include an operon adjacent to ColR/S that includes *orf222* (a putative methyltransferase) and *wapQ* (*a* putative heptose kinase), which are predicted to modify bacterial lipopolysaccharide (LPS) [20]. Additionally, ColR/S in *P. putida* regulates expression of a phosphoethanolamine (pEtN) transferase *eptA*, which adds a pEtN to LPS [21]. We found that all three genes are conserved in both protective and non-protective *Pseudomonas* strains (Fig 4A). We generated deletion mutants in *Pseudomonas* sp. WCS365 *eptA, orf222* and *wapQ* and tested the mutants for protection against N2C3. We found that similar to the Δ*colR* mutant, the WCS365 Δ*wapQ* mutant failed to protect against N2C3 (Figure B). In contrast the Δ*orf222* and Δ*eptA* mutants retained their ability to protect against N2C3. These results indicate that a subset of ColR-dependent genes are required for protection against N2C3.

**Figure 4.**
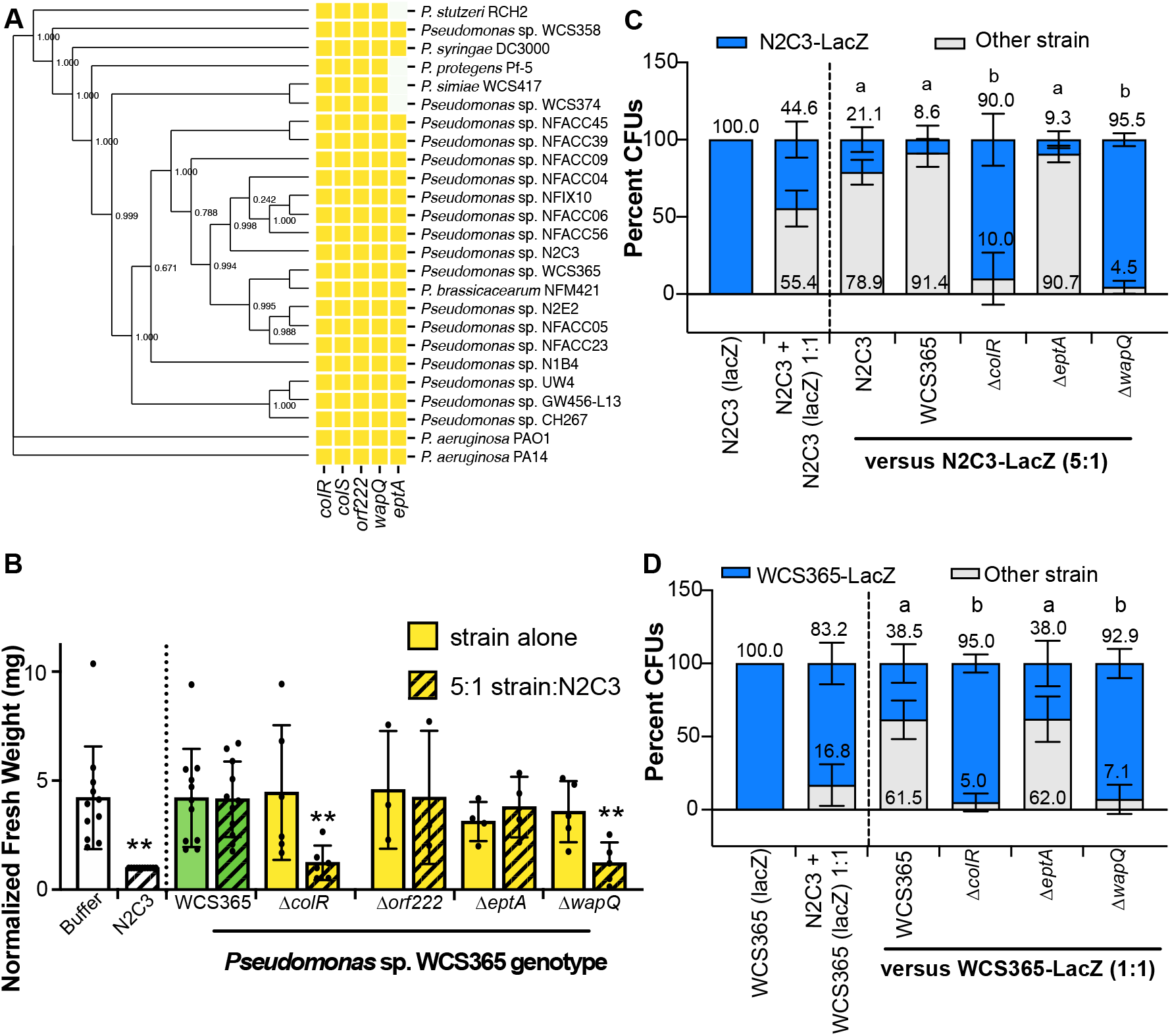
Protection against N2C3 is dependent on *colR* and *wapQ*. (A) The two-component system ColR/S and three previously identified genes whose expression is ColR/S dependent are largely conserved across both protective and non-protective *Pseudomonas* strains. (B) A deletion mutant in *Pseudomonas* sp. WCS365 *wapQ* was unable to protect against N2C3 when co-inoculated at a 5:1 ratio. Data was normalized by dividing by average fresh weight of N2C3-inoculated control plants. **p<0.05 by one-way ANOVA and Dunnett’s post-hoc test. Data for WCS365 strains inoculated alone were excluded from statistical analyses. (C) Deletion of the ColR-regulated gene, *wapQ*, but not *eptA*, results in significantly decreased fitness in competition with N2C3 and (D) WCS365. When competed against N2C3-lacZ in a 5:1 ratio and WCS365-lacZ at a 1:1 ratio, non-protective WCS365 deletion mutants (WCS365Δ*colR* and WCS365Δ*wapQ*) were significantly outcompeted.

To test whether Δ*colR* and Δ*wapQ* mutants had competition defects in the rhizosphere, we competed them against N2C3-lacZ and WCS365-lacZ. We found that protective strains WCS365 and WCS365Δ*eptA* maintained their inoculated ratios in the rhizosphere in competition with either N2C3 or WCS365 (Fig 4C-D), whereas the non-protective strains WCS365 Δ*colR* and WCS365 Δ*wapQ* were significantly out-competed by N2C3 (Fig 4C) or WCS365 (Fig 4D). These data support the observation amongst diverse *Pseudomonas* strains that protection correlates with rhizosphere colonization. Furthermore, only a subset of *colR*-dependent genes, which have rhizosphere fitness defects, were found to be necessary for protection.

### Commensal *Pseudomonas* can protect against an agricultural pathogen

To determine how broadly relevant the system we developed is to identify members of the microbiome that can protect against *Pseudomonas* pathogens, we tested whether *Pseudomonas* sp. WCS365 was able to protect against *P. fuscovaginae* strain SE-1, the causal agent of rice brown sheath rot, which uses syringomycin and syringopeptin as virulence factors [22]. We found that WCS365 was able to protect against *P. fuscovaginae* strains SE-1 (Fig 5). Interestingly, although WCS365 rescued fresh weight, shoot growth, and lateral root formation in the presence of SE-1, it did not fully rescue primary root stunting. Collectively these data indicate that the described gnotobiotic assay may be able to broadly identify commensal strains and mechanisms that protect from agriculturally important pathogens.

**Figure 5.**
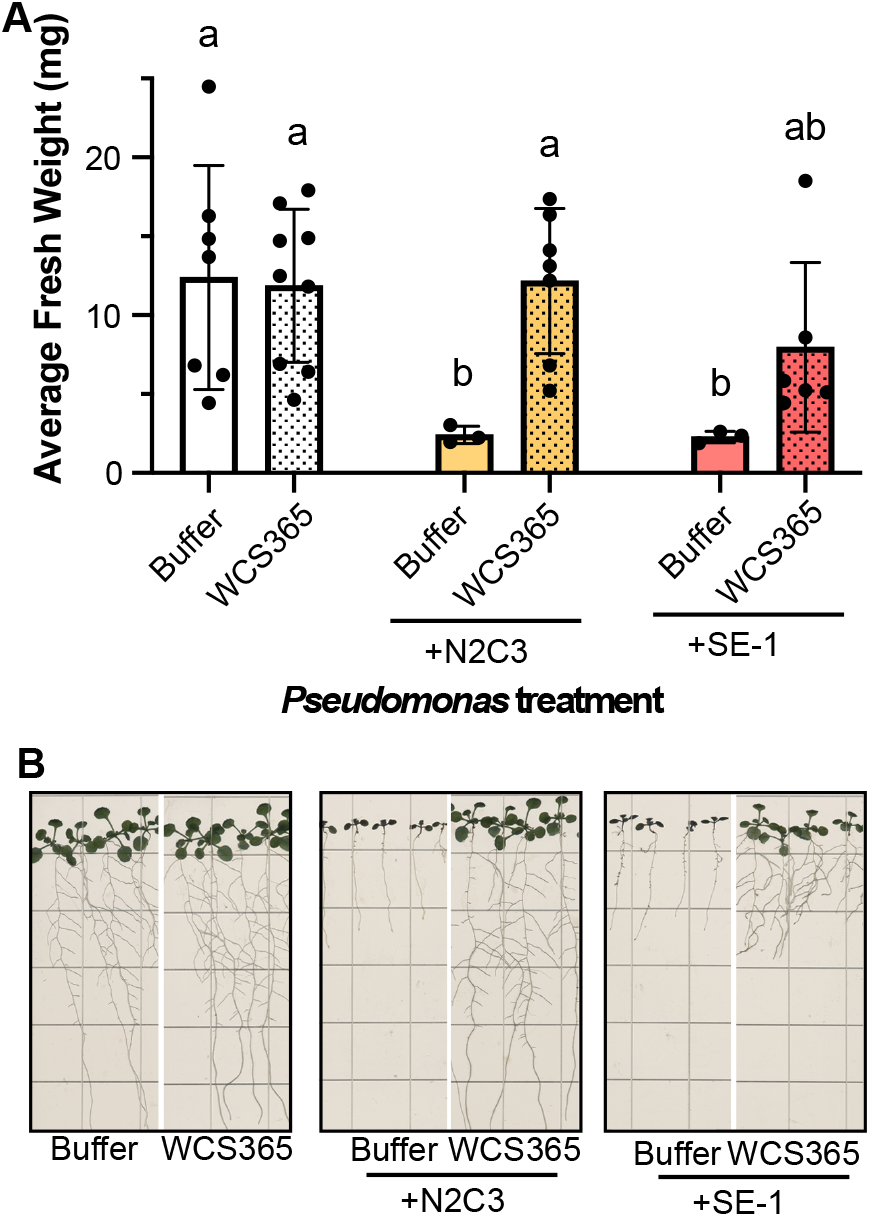
WCS365 protects against the agronomically important *Pseudomonas fuscovaginae* pathogen SE-1. (A) WCS365 protects against SE-1, the causal agent of bacterial sheath brown rot of rice when co-inoculated onto *Arabidopsis* roots at a 5:1 ratio. Letters indicate p<0.05 determined by one-way ANOVA and Tukey HSD. (B) Representative images of data quantified in (A).

## Discussion

Using a reductionist system consisting of *Arabidopsis*, an opportunistic *Pseudomonas* pathogen, and commensal *Pseudomonas*, we identified strains and genes required for members of the rhizosphere microbiome to protect against pathogens. We found that the ability to protect against pathogenesis by *Pseudomonas* sp. N2C3 was almost uniquely present within the BCM clade, indicating that strains closely related to N2C3 could protect against its pathogenesis, while more distantly related *Pseudomonas* strains could not. However, unlike many traits that show phylogenetic signatures within the genus *Pseudomonas* [9,10,14], we were unable to identify components of the *Pseudomonas* accessory genome that correlate with protection. As a result, this work implicates variation in regulation of components of the core genome in protection against an opportunistic pathogen.

A subset of previously identified colonization genes, *colR* and *wapQ*, were found to be necessary for protection by WCS365. ColR has been previously reported to impact rhizosphere colonization, by affecting membrane permeability via *wapQ* [20]. As ColR has been identified in multiple screens for rhizosphere fitness determinants, this may indicate that the *ΔcolR* mutant has the most severe colonization/fitness defect of those tested [18,19,23]. It may be that closely-related strains have similar metabolic potential as N2C3 and so as long as they can colonize at high enough levels, they can restrict pathogen growth and suppress disease. However, as not all colonization factors or *colR*-dependent genes were necessary for WCS365 to protect against N2C3, a specific *colR*-dependent colonization mechanism may be required for protection.

We found that the protective strain *Pseudomonas* sp. WCS365 was unable to inhibit N2C3 growth *in vitro* suggesting that inhibition requires the presence of a plant. Protective strains co-inoculated with N2C3 tend to maintain approximately the same co-inoculation ratio 7 days after inoculation. This demonstrates that N2C3 is not excluded by the protective strains, but rather persists in the rhizosphere as a commensal member of the microbiome. This suggests that N2C3 transitions between pathogenic and non-pathogenic lifestyles, depending on the presence of specific microbiota, which is characteristic of opportunistic pathogens. Closely-related *Pseudomonas* strains with markers of both commensal and pathogenic lifestyles ([9,11] and NFACC strains described in this study) have been identified within the microbiome of the same plant suggesting competition may be ecologically relevant. This also suggests that a lack of soil diversity could lead to increased opportunities for pathogens to attack. As we found that *Pseudomonas* sp. WCS365 was also able to protect against *P. fuscovaginae* SE-1, this system may be broadly relevant for identifying strains and mechanisms to control agriculturally important pathogens.

## Materials and Methods

### Bacterial cultures and growth media

Strains used in this work are summarized in Table 1. All *Pseudomonas* spp. were routinely cultured on King’s B or LB agar plates, and incubated at 28°C. *Escherichia coli* and *Chromobacterium violaceum* CV026 were cultured on LB agar plates, and incubated at 37°C and 28°C, respectively. Overnight cultures of *Pseudomonas* spp. and *C. violaceum* CV026 were prepared in 5 mL of LB, grown at 29°C, and shaken at 180 rpm. Overnight cultures of *E. coli* were prepared in 5 mL of LB, grown at 37°C, and shaken at 180 rpm. When required, growth media were supplemented with 25 μg/mL gentamycin, 10 μg/mL nalidixic acid, 10% sucrose, or 0.2 mg/mL X-gal.

NFACC and NFIX strains used in this study are a part of a bacterial collection isolated from the roots of Switchgrass (*Panicum virgatum*) growing in Tallgrass Prairie Preserve, Oklahoma, USA. Approximately 10 pieces of healthy roots (1 cm long) were surface sterilized and crushed with a pestle to extract sap. Diluted sap were plated on 869 media. After 3-4 days of incubation, 3-4 colonies of different shapes were purely isolated from each plant. Their genomes are available at https://genome.jgi.doe.gov/portal/Comgentwoseasons/Comgentwoseasons.info.html

### Plant growth conditions

*Arabidopsis thaliana* Col-0 seeds were sterilized by submersion in 70% ethanol for 2-3 minutes, then 10% bleach for 1-2 minutes. The seeds were rinsed three times using sterile deionized water, then submerged into 0.1% phytoagar. The seeds were stored in the dark at 4°C for at least 2 days prior to sowing. The plants were grown in a growth room using a 16h light/8h dark cycle, under 100 μM fluorescent white light.

### Axenic root inoculation assays

Sterilized seeds were planted onto square plates containing 0.5X Murashige and Skoog (MS) media, with 0.5 g/L 2-(N-morpholino)ethanesulfonic acid (MES) buffer, 2% sucrose, and 1% phytoagar. The plates were sealed using Micropore tape and placed upright, allowing seedlings to germinate along the surface of the media. After 6 days, the seedlings were carefully transferred onto square plates containing 0.5X MS media, with 0.5 g/L MES buffer, no sucrose, and 1% phytoagar, before being resealed and returned to the growth room. On day 7, seedlings were inoculated along their roots with 5 μL of bacterial treatments, prepared as described below. 5 seedlings were inoculated per treatment.

Bacteria were prepared by inoculating single colonies into 5 mL of LB, to grow overnight cultures. 1 mL of overnight culture was centrifuged for 1 min at 10000 rcf, and the pelleted cells were resuspended in 1 mL of 10 mM MgSO_4_. The resuspended cells were serially diluted to an approximate OD_600_ of 0.001 using 10 mM MgSO_4_. Bacterial mixtures were prepared at 5:1, 1:1, and 1:5 ratios of test strain:N2C3 where “1” is 10 μL and “5” is 50 μL. For example, a 5:1 treatment of WCS365:N2C3 would contain 50 μL of WCS365 and 10 μL of N2C3.

After bacterial inoculation, the plates were resealed and the plants were grown vertically for 7 days in the growth room. Images of the plates were then scanned, and seedlings from the same treatment were pooled for fresh weight measurements.

### Rhizosphere CFU counts

N2C3-lacZ and WCS365-lacZ strains were generated through biparental mating of the N2C3 or WCS365 with *E. coli* WM3064 or CC18 containing *pMini-Tn5-lacZ*, respectively. The relative fitness ratio (ω) of lacZ strains was calculated for the transconjugant versus the parental strain in planta from 7-14 days in planta [24]. For N2C3-lacZ vs N2C3 ω = 0.993 +/− 0.024 and for WCS365-lacZ vs WCS365 ω = 1.012 +/− 0.038 indicating that the LacZ transposon insertion did not result in a significant change in fitness in either strain. For quantification of rhizosphere colony-forming unit (CFU) counts, either N2C3 or WCS365 strains containing a *pMini-Tn5-lacZ* insertion were used to co-inoculate 7-day-old *Arabidopsis* seedlings, as described in the Axenic Root Inoculation Assay protocol.

After 7 days, seedlings were pooled and sterilely transferred into a 2 mL microcentrifuge tube containing a metal bead and 500 μL of 10 mM MgSO_4_. Fresh weights were measured for each treatment. Plant cells were homogenized using a Qiagen TissueLyser II at 30 Hz for 2 min. Tissue lysate was serially diluted and plated onto LB agar supplemented with 0.2 mg/mL 5-Bromo-4-Chloro-3-Indolyl β-D-Galactopyranoside (X-gal). Plates were incubated at 28°C for 2 days. Blue and white colonies were counted, to calculate the ratio of blue LacZ-containing cells and white co-inoculated cells in the rhizosphere.

### AHL biosensor assays

Overnight cultures of *Pseudomonas* sp. WCS365, *Pseudomonas* sp. N2C3, and *C. violaceum* CV026 were resuspended then diluted in 10 mM MgSO_4_, to an OD_600nm_ of 2. Bacterial treatments were prepared by mixing 5:1, 1:1, and 1:5 ratios of WCS365:N2C3. Using an inoculation loop, bacterial treatments were streaked along an LB agar plate. CV026 was streaked beside each bacterial treatment. Plates were incubated at 28°C and imaged after 1 day.

### Comparative genomics using PyParanoid

Comparative genomics was performed using the PyParanoid pipeline described previously[9,10]. Briefly, the PyParanoid pipeline was previously used to create a database using 3886 *Pseudomonas* genomes, identifying 24066 homologous protein families (“gene groups”) that covered 94.2% of the generated *Pseudomonas* pangenome. The presence or absence of gene groups were compared in strains with protection phenotypes determined via axenic root inoculation assays.

### Phylogenetic trees

Phylogenetic trees were generated using the PyParanoid pipeline and FastTree 2 as described previously [9,10]. Briefly, an alignment of 122 single-copy genes conserved within the *Pseudomonas* genus was created previously. FastTree 2 was used to generate a phylogenetic tree from this alignment, by randomly sampling 5000 amino acid residues without replacement. Strains were then subset from this alignment to create phylogenetic trees, using FastTree 2.

### Generating gene deletions

Gene deletions in WCS365 were created using a two-step allelic exchange method described previously [25]. For each target gene deletion, 700-900 bp regions immediately flanking upstream and downstream of the target region were amplified then using WCS365 genomic DNA as a template, and the primers listed in Supplementary Table 1. The amplified flanking regions were then joined using overlap PCR. The overlap PCR product was digested and ligated into the pEXG2 suicide vector, and transformed into competent *Escherichia coli* DH5α cells via heat-shock. Correct insertions were confirmed by colony PCR and Sanger sequencing, before transforming the plasmid into *E. coli* SM10λpir. Deletion plasmids were conjugated into WCS365 through biparental mating. Transconjugants that underwent a single crossover event allowing for site-specific chromosomal integration were selected for using LB agar supplemented with 25 μg/mL of gentamycin and 15 μg/mL of nalidixic acid. After restreaking and validating antibiotic resistance, transconjugants were grown overnight in no-salt LB at 37°C, and cells that underwent a double crossover to excise the plasmid backbone were selected by plating on LB agar supplemented with 10% sucrose. Successful gene deletions were confirmed using PCR, gel electrophoresis, and Sanger sequencing.

### Cross-streak assays

Bacteria were grown overnight in LB, at 28°C. Using a sterile inoculation loop, the first bacterial strain was streaked onto LB plates, directly from the overnight culture. The plate was allowed to dry, before streaking the second strain was streaked perpendicular to the first. The plate was incubated overnight at 28°C.

### Statistical analyses

All statistical analyses were performed using Prism 9 software. In general, competition assays were analyzed by one-way ANOVA and Tukey’s HSD test and LacZ assays were analyzed using two-way ANOVA and Tukey’s HSD test.

## Acknowledgements

We thank Dr. Kelly Craven for sharing NFACC and NFIX strains. This work was supported by an NSERC Discovery Grant (NSERC-RGPIN-2016-04121) and Weston Seeding Food Innovation grants awarded to CHH. NRW was supported by an NSERC CGS-M and an NSERC training CREATE grant and CLW was supported by an NSERC CGS-D award. RAM was a Simons Foundation Fellow of the Life Sciences Research Foundation. The computational research was carried out with support provided by WestGrid and Compute Canada.

