## Supplementary Data for "Commensal *Pseudomonas fluorescens* protect *Arabidopsis* from closely-related *Pseudomonas* pathogens in a colonization-dependent manner"

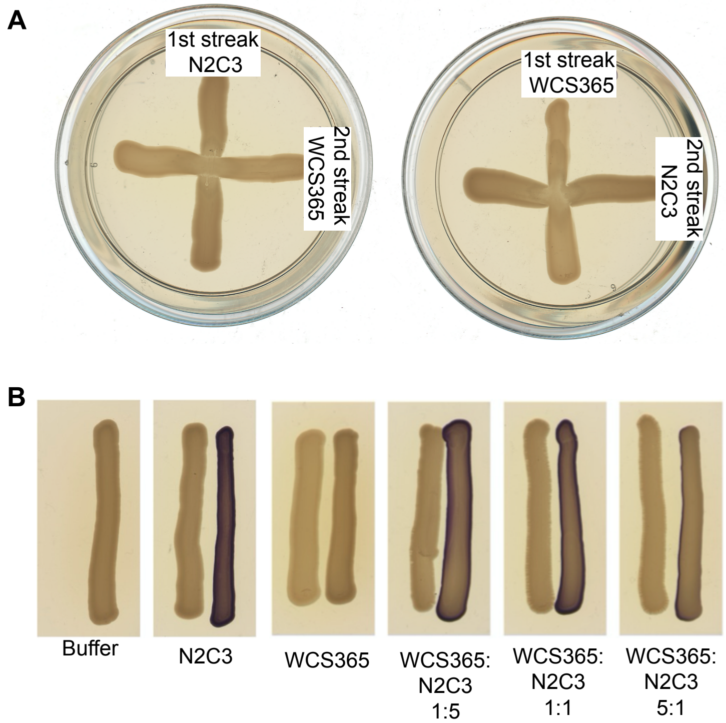


**Figure S1.** ***Pseudomonas* sp. WCS365 does not inhibit growth or quorum signaling of N2C3 *in vitro*.** (A) Overnight cultures of WCS365 and N2C3, grown in LB, were streaked onto LB agar. The first strain was streaked vertically, and the second strain was streaked horizontally at a 90° angle. Neither WCS365 nor N2C3 created a zone of inhibition that prevented the growth of the strain that was streaked second. (B) Although N2C3 virulence relies on quorum signalling genes present in a pathogenicity island in the BCM clade, the protective WCS365 strain does not protect through quorum quenching. N2C3 AHL production is detected by the production of violacein, a visible purple pigment, by *C. violaceum* CV026. Each panel contains two streaks. On the left is the bacterial mixture described below each panel. On the right is the CV026 biosensor, which produces violacein in the presence of C4-C8 AHL molecules.

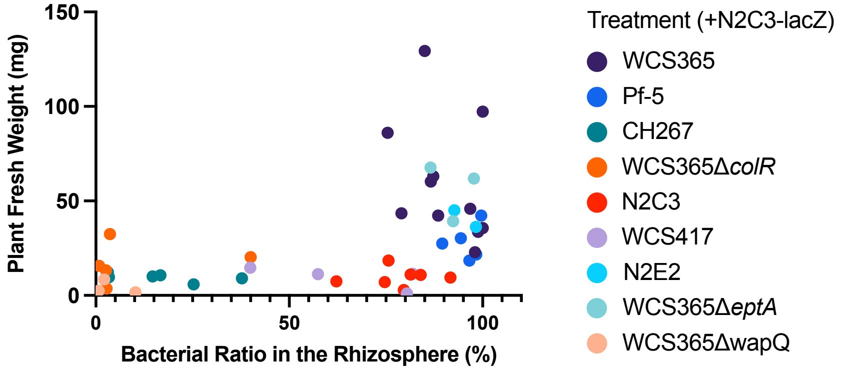


**Figure S2. Protection against *Pseudomonas* sp. N2C3 is rhizosphere colonization dependent.** Correlation between the bacterial fitness in the rhizosphere and plant fresh weight. Data points represent the averaged fresh weights for individual biological replicates, color coded by treatment from the data shown in Figure 3A-B and Figure 4C. Each strain was co-inoculated with N2C3-lacZ at a 5:1 ratio.

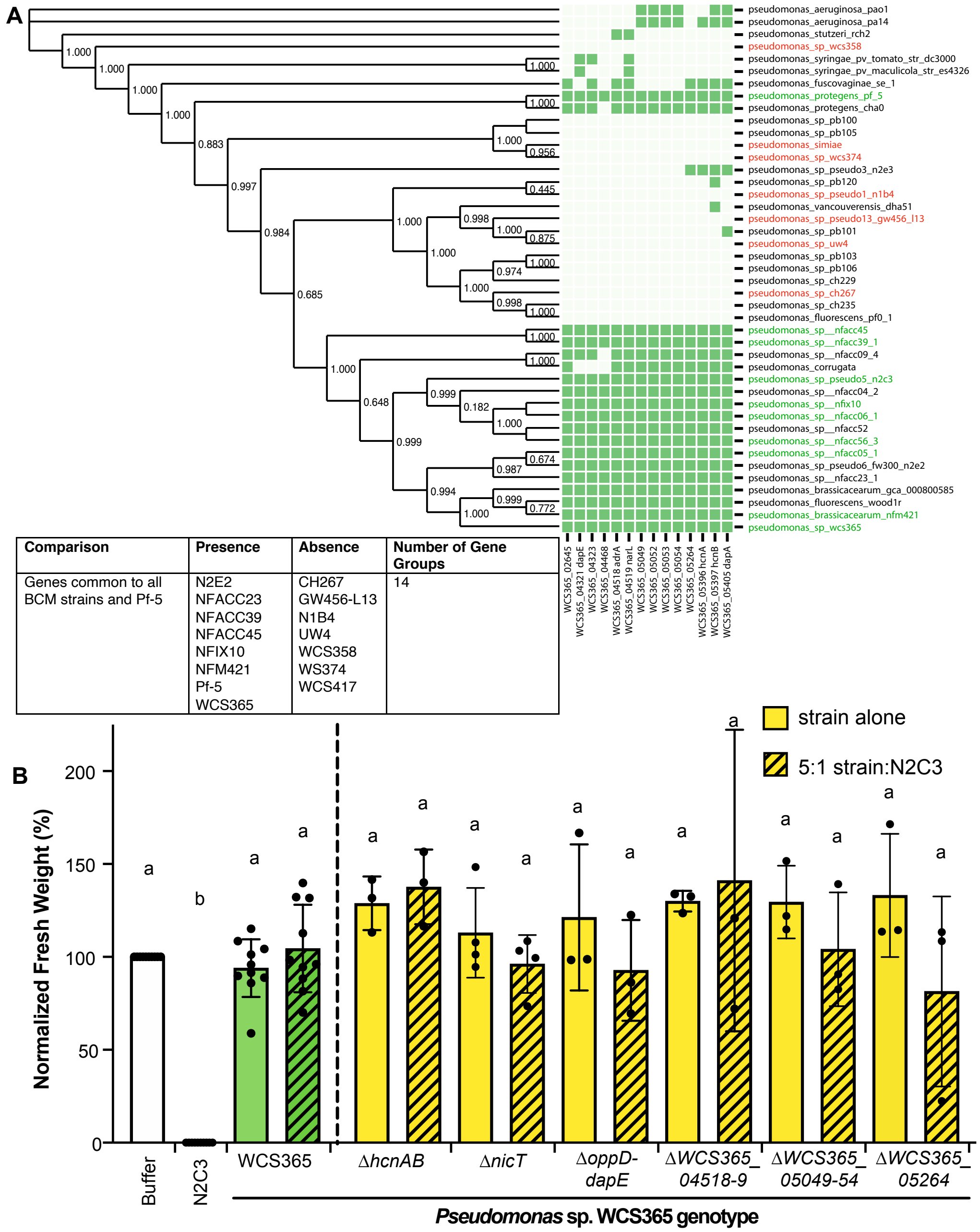


| Comparison | Presence | Absence | Number of Gene Groups |
|---|---|---|---|
| Genes common to all BCM strains and Pf-5 | N2E2<br>NFACC23<br>NFACC39<br>NFACC45<br>NFIX10<br>NFM421<br>Pf-5<br>WCS365 | CH267<br>GW456-L13<br>N1B4<br>UW4<br>WCS358<br>WS374<br>WCS417 | 14 |

**Figure S3. Comparative genomics analyses performed to identify genes common to BCM strains and Pf-5** (A) To test whether Pf-5 obtained genes responsible for protection in the BCM clade through horizontal gene transfer, we identified 14 genes that were present in 7 BCM strains and Pf-5 (green labels), but absent in 7 non-protective strains outside of the BCM clade (red labels). Both protective and non-protective BCM strains were included in this comparison to capture genes that may be differentially expressed. Gene groups were labelled by their locus tag in WCS365. (B) 12/14 of the genes identified were deleted in WCS365, but none of them had an impact on its ability to protect against N2C3. Data was normalized by dividing by average fresh weight of N2C3-inoculated plants. Statistics were calculated using a one-way ANOVA and Tukey's HSD. Mean +/- SD is plotted.

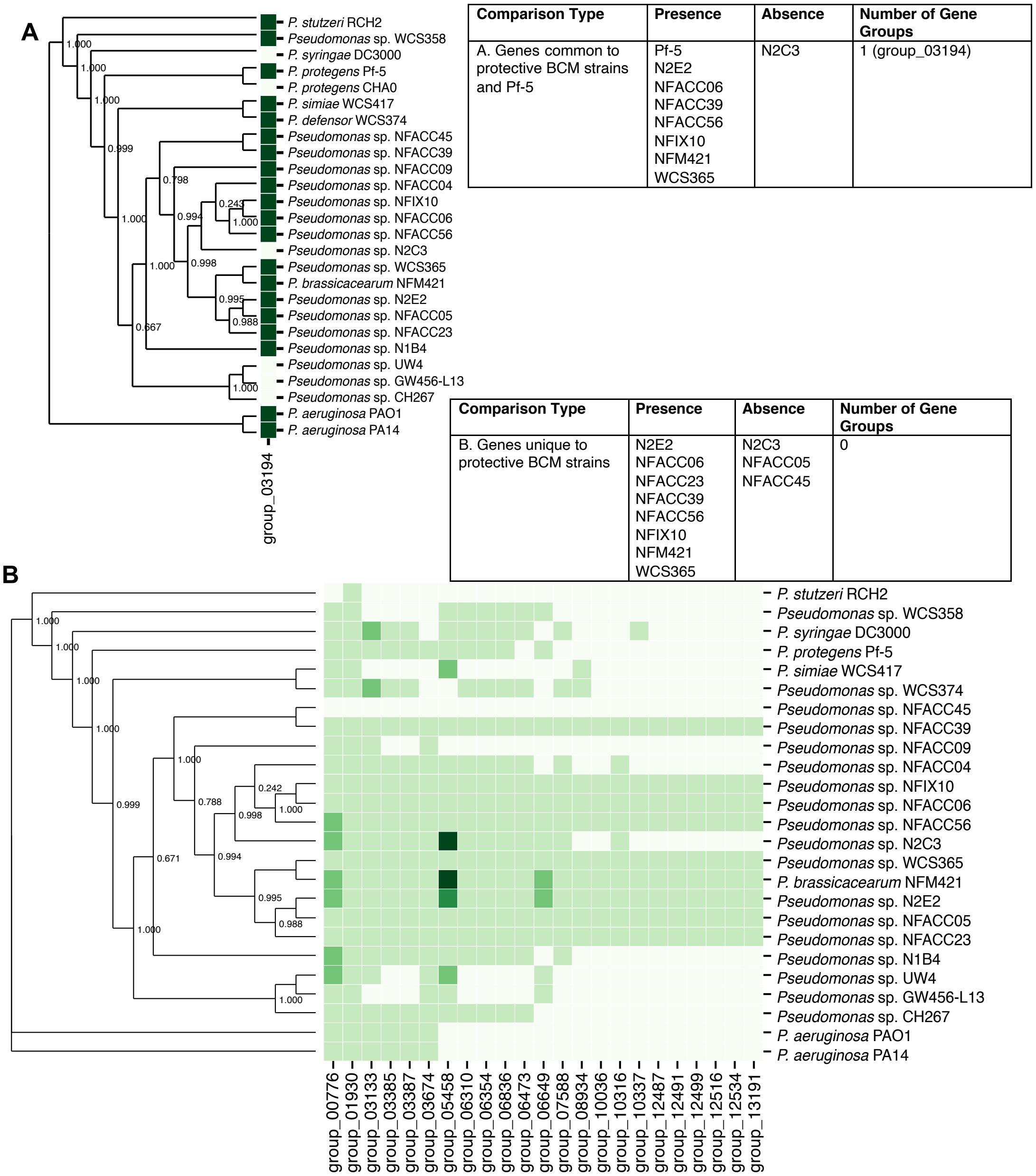


| Comparison Type | Presence | Absence | Number of Gene Groups |
|---|---|---|---|
| A. Genes common to protective BCM strains and Pf-5 | Pf-5<br>N2E2<br>NFACC06<br>NFACC39<br>NFACC56<br>NFIX10<br>NFM421<br>WCS365 | N2C3 | 1 (group_03194) |

| Comparison Type | Presence | Absence | Number of Gene Groups |
|---|---|---|---|
| B. Genes unique to protective BCM strains | N2E2<br>NFACC06<br>NFACC23<br>NFACC39<br>NFACC56<br>NFIX10<br>NFM421<br>WCS365 | N2C3<br>NFACC05<br>NFACC45 | 0 |

**Figure S4. Comparative genomics analyses performed to identify genes unique to protective strains.** (A) Using PyParanoid, we searched for genes present in seven protective BCM strains (NFM421, N2E2, WCS365, NFIX10, NFACC06, NFACC39, and NFACC56), the protective non-BCM strain Pf-5, but absent in N2C3. This approach yielded only 1 unique gene (group_03914), encoding a CinA family protein. However, this gene was subsequently found to be present in 9 non-protective strains (NFACC05, NFACC45, N1B4, PA14, PAO1, RCH2, WCS358, WCS374, and WCS417) and absent in the protective strain CHA0, indicating that it is unlikely to underlie protection. (B) A comparative genomics analysis between strains within the BCM clade yielded no genes unique to protective BCM strains. We compared 8 protective BCM strains (NFM421, N2E2, WCS365, NFIX10, NFACC23, NFAC45, NFACC39, NFACC06, and NFACC56) with 3 non-protective BCM strains (NFACC45, NFACC05, and N2C3). Using this approach, we identified no genes present in the protective BCM strains, but absent in all 3 non-protective BCM strains.

A

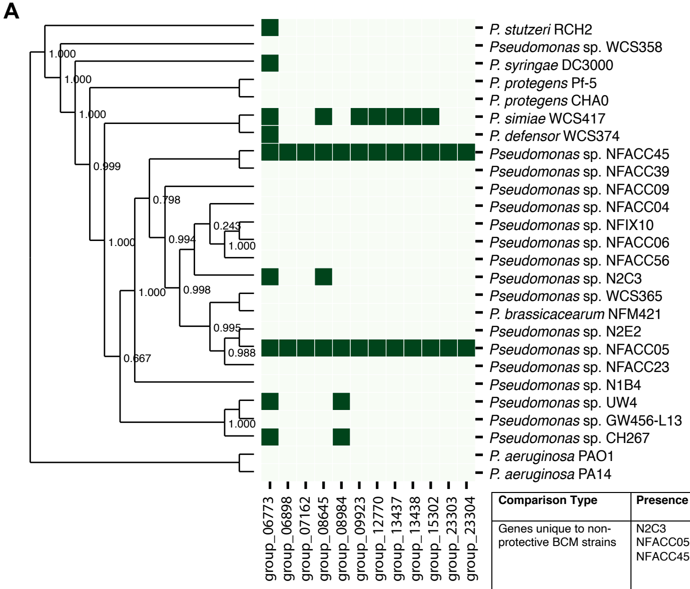


| Comparison Type | Presence | Absence | Number of Gene Groups |
|---|---|---|---|
| Genes unique to non-protective BCM strains | N2C3<br>NFACC05<br>NFACC45 | N2E2<br>NFACC06<br>NFACC23<br>NFACC39<br>NFACC56<br>NFIX10<br>NFM421<br>WCS365 | 1 (group_06773) |

B

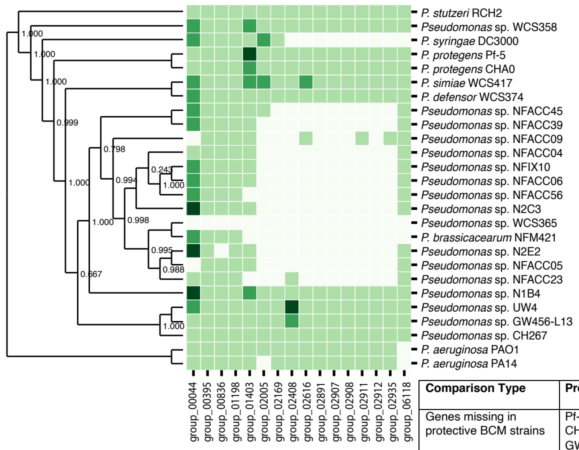


| Comparison Type | Presence | Absence | Number of Gene Groups |
|---|---|---|---|
| Genes missing in protective BCM strains | Pf-5<br>CH267<br>GW456-L13<br>N1B4<br>RCH2<br>UW4<br>WCS358<br>WCS374<br>WCS417 | N2E2<br>NFACC06<br>NFACC39<br>NFACC56<br>NFIX10<br>NFM421<br>WCS365 | 0 |

**Figure S5. Comparative genomics analyses to identify genes absent from protective BCM strains.** (A) To identify genes that are unique to non-protective BCM strains, we searched for genes that were present in 3 non-protective BCM strains (NFACC45, NFACC05, and N2C3), but absent in 8 protective BCM strains (NFM421, N2E2, WCS365, NFIX10, NFACC23, NFAC45, NFACC39, NFACC06, and NFACC56). Using this approach, we identified 1 gene group encoding an aldo/keto reductase (group_06773) that was unique to non-protective BCM strains. However, not all non-protective strains outside of the BCM clade possessed this gene group, indicating that the absence of this gene was unlikely to be responsible for protection in the BCM clade. (B) To identify genes whose absence may be responsible for a BCM clade-specific mechanism of protection, we searched for genes that were present in 9 non-BCM strains (Pf-5, CH267, GW456-L13, N1B4, RCH2, UW4, WCS358, WCS374, and WCS417) but absent in 7 protective BCM strains (N2E2, NFACC06, NFACC39, NFACC56, NFIX10, NFM421, and WCS365). However, we were unable to identify any genes that met these criteria.

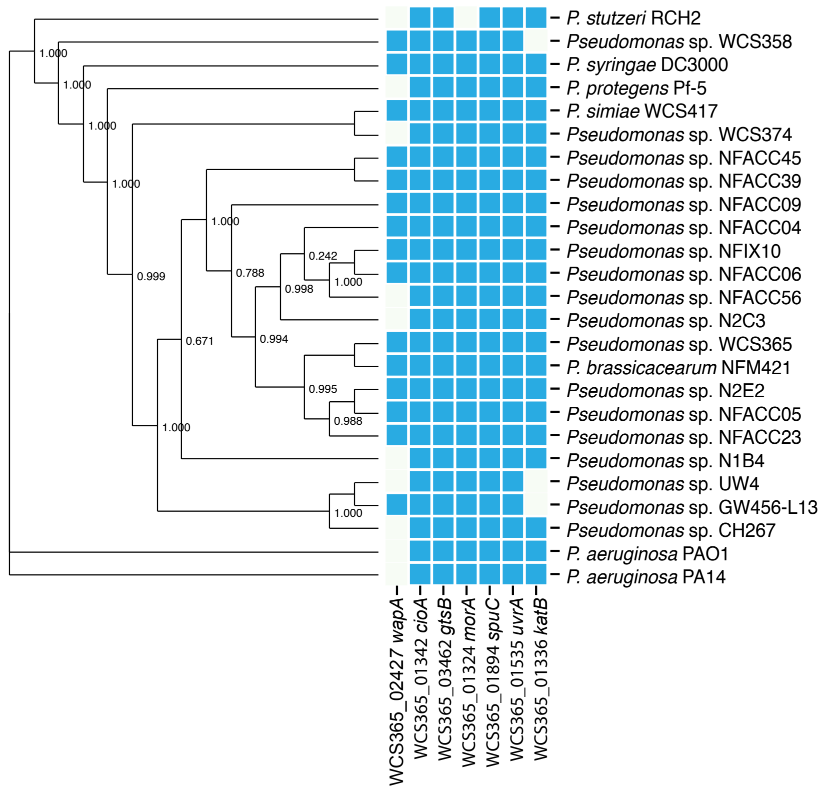


**Figure S6. Distribution of colonization genes across *Pseudomonas* sp.** Previously identified genes involved in colonization and rhizosphere fitness whose expression is ColR/S dependent are largely conserved across both protective and non-protective *Pseudomonas* strains.

**Table S1. Primers used to generate WCS365 and N2E2 mutants**

| Deletion Strain | Primer Type | Primer Sequence (5' ⟶ 3') | RE Site |
|---|---|---|---|
| WCS365 Δ*hrcC* | Upstream forward | GNNAAGCTTTGCCTGTGAACACCCCGGAA | HindIII |
| | Upstream reverse | ACCATCGACTGCATTTATATAGTGTACTTTGCCTGCGGTCAT | |
| | Downstream forward | TATATAAATGCAGTCGATGGTGTCATGAAACAACCCACGCG | |
| | Downstream reverse | GNNAAGCTTGAGCTATGGAGGATGGCGAC | HindIII |
| WCS365 ΔDAPG | Upstream forward | GNNAAGCTTGCAATTCACGCGACGTATCC | HindIII |
| | Upstream reverse | ACCATCGACTGCATTTATATACGCGCCATAGCTCACAATTC | |
| | Downstream forward | TATATAAATGCAGTCGATGGTGGCTGTATTGACCGGCCCCTG | |
| | Downstream reverse | GNNAAGCTTGACCAGTGAGAGTGTCGAGC | HindIII |
| WCS365 Δ*eptA* | Upstream forward | ATATGAATTCGCATGCACATGCATGTCAGCC | EcoRI |
| | Upstream reverse | GACAGGTTAAACAACGCTGCGTGACGGCTTCCTGAAAGTTGCTC | |
| | Downstream forward | GAGCAACTTTCAGGAAGCCGTCAACGCAGCGTTGTTTAACCTGTC | |
| | Downstream reverse | ATATGGATCCGCCTAAGAGAAATCGCCTGCG | BamHI |
| | Upstream confirmation | GCAGGCCTGCTTCATGGC | |
| | Downstream confirmation | CTTGAAGAATGGCGCGCAG | |
| WCS365 Δ*wapQ* | Upstream forward | ATATGAATTCCCCATCAATGCCCGACCC | EcoRI |
| | Upstream reverse | CTGCCAGGGCTCCGACTAAAATCGTTGATGGTTCTGCTGCC | |
| | Downstream forward | AATCGTTGATGGTTCTGCTGCCTTTAGTCGGAGCCCTGGCAG | |
| | Downstream reverse | ATATGGATCCGCGTCGACTCGTCTGC | BamHI |
| | Upstream confirmation | ACGGTTGCCGTTTCGAGG | |
| | Downstream confirmation | GCCGTGGAACAAGCCGC | |
| WCS365 Δ*orf222* | Upstream forward | ATATGAATTCTTGGCGACCTCCTGCG | EcoRI |
| | Upstream reverse | CCTGAATACGAAACCCTGCCTGTGATGGGGGCCTCCATGTC | |
| | Downstream forward | GACATGGAGGCCCCCATCACAGGCAGGGTTTCGTATTCAGG | |
| | Downstream reverse | ATATGGATCCCGACGAGTGGCGCCG | BamHI |
| | Upstream confirmation | CCGTGGCGTTCGACGC | |
| | Downstream confirmation | AACTGCCAGGGCTCCG | |

**Table S2. Primers used to generate WCS365 mutants from comparative genomics analyses**

| **Deletion Strain** | **Primer Type** | **Primer Sequence (5' ⟶ 3')** | **RE Site** |
|---|---|---|---|
| WCS365 Δ*nicT* | Upstream forward | CATCAAGCTTCAGCAGCAGCGCTATGACTT | HindIII |
| | Upstream reverse | AGGATCAGTCCACCGCTGACGCAACTGCTCGGAGATGGTT | |
| | Downstream forward | AACCATCTCCGAGCAGTTGCGTCAGCGGTGGACTGATCCT | |
| | Downstream reverse | CATCGGATCCAATTGCCGGTAGACCTGTGC | BamHI |
| | Upstream confirmation | ATCCTGCAGATCAAGCAGCG | |
| | Downstream confirmation | TGCGCGATCTCTTCAGTGTG | |
| WCS365 Δ*hcnAB* | Upstream forward | CATCAAGCTTGCGAAGCCTGCGAACTGATC | HindIII |
| | Upstream reverse | GCTATGGCGCAGTGTCAGTCAAACAAGTCAGGCATGGGCC | |
| | Downstream forward | GGCCCATGCCTGACTTGTTTGACTGACACTGCGCCATAGC | |
| | Downstream reverse | CATCGGATCCCGCCCCATCAGGGAACAAGA | BamHI |
| | Upstream confirmation | TGATGGATTGGCCCTGTGCC | |
| | Downstream confirmation | GCCGATGATGTCATCGAGCT | |
| WCS365 Δ*oppD-dapE* | Upstream forward | tcatGGATCCCCGATGTGCTCAACGCACTG | BamHI |
| | Upstream reverse | CCCGCCTTGTCATAGCTGTTGCACCAGAATCATCAGGGCG | |
| | Downstream forward | CGCCCTGATGATTCTGGTGCAACAGCTATGACAAGGCGGG | |
| | Downstream reverse | catgAAGCTTGTTCGCTGTGACAGGCTTCC | HindIII |
| | Upstream confirmation | CGCGTGTTGTTCAAGCATGC | |
| | Downstream confirmation | ACGCTCGGTATGAACCTTGC | |
| WCS365 Δ04518-9 | Upstream forward | tcatGGATCCCATCCTTGGCGATCCGGTTT | BamHI |
| | Upstream reverse | CCAGCCACCCTCAATGTTGCCCATTGGCGTTCAATGGCAG | |
| | Downstream forward | CTGCCATTGAACGCCAATGGGCAACATTGAGGGTGGCTGG | |
| | Downstream reverse | gcatAAGCTTGAAAGTGGCTGGGACTTCAGC | HindIII |
| | Upstream confirmation | GTCGAGCCGTCGACTGAAAC | |
| | Downstream confirmation | CCTCGACAGGAATCCTGGCT | |
| WCS365 Δ05049-54 | Upstream forward | tcatGGATCCCGATTATCACTGGCCGCGTC | BamHI |
| | Upstream reverse | AAACATCCGGATCAACGCCCATGGTGATGTTCTTGCCGCT | |
| | Downstream forward | AGCGGCAAGAACATCACCATGGGCGTTGATCCGGATGTTT | |

| | | | |
|---|---|---|---|
| | Downstream reverse | gcatAAGCTTTAGCTCGCCTCTGAAGAGGC | HindIII |
| | Upstream confirmation | CTGGGATCGCCATGACCAGT | |
| | Downstream confirmation | AGCGTGATCTGGATATCGGCT | |
| WCS365 Δ*05264* | Upstream forward | gcatAAGCTTGACAATCTTGGCGCGAGTCC | HindIII |
| | Upstream reverse | CTAGTCCGGCGATGCTATCCCAGGACACGCCAGTCTGTTG | |
| | Downstream forward | CAACAGACTGGCGTGTCCTGGGATAGCATCGCCGGACTAG | |
| | Downstream reverse | tcatGGATCCCGGCATCGAAGTAGGCGTAG | BamHI |
| | Upstream confirmation | CTCAATCGGATGGGCCATCAA | |
| | Downstream confirmation | CGACGGCTTCGTCGTTGATC | |
